# Divergent transcriptomic transition programs in Alzheimer disease: immune priming and synaptic collapse in females versus organelle membrane bifurcation in males

**DOI:** 10.64898/2026.07.09.735789

**Authors:** Moonseok Choi, Sarah Bauermeister, Do-Geun Kim

## Abstract

Alzheimer’s disease (AD) disproportionately affects women, who account for approximately two-thirds of prevalent cases. Despite decades of research, the mechanistic basis for this profound sex disparity remains poorly resolved. Prior transcriptomic studies have predominantly used pooled or female-enriched cohorts, obscuring whether the transition from normal cognition (NCI) to AD follows a common molecular program across sexes or reflects fundamentally divergent biology. Here, we analyzed bulk RNA-sequencing data from the dorsolateral prefrontal cortex of 624 individuals (401 females, 223 males) in the ROSMAP cohort using a transition-aware framework that separates variance instability along the disease axis from mean expression changes. We demonstrate that female and male brains exhibit structurally distinct transcriptomic transition programs. Females display a sequential, multi-tier architecture: interferon and immune variance priming (628 genes) is detectable early at the NCI-to-mild cognitive impairment (MCI) interval, which structurally precedes a massive mean-level synaptic and neuropeptide loss (8,935 genes) in AD. The female NCI-to-MCI interval alone produces 1,249 variance bifurcation events entirely absent in males. Conversely, males exhibit no early immune priming or powered mean-level changes. Instead, they collapse into a single large variance bifurcation pool (8,237 genes) heavily enriched for system-wide organelle membrane fusion, post-translational modification, and intracellular trafficking. These findings reveal that the AD transcriptomic transition is not a unitary program quantitatively modified by sex, but two distinct biological trajectories. This fundamental divergence motivates sex-stratified mechanistic models and independent biomarker development for each sex, cautioning against analytical pooling in transition-stage cohorts.

## Main

Alzheimer’s disease (AD) is the most prevalent cause of dementia globally, yet its clinical burden is profoundly asymmetrical, disproportionately affecting women, who account for approximately two-thirds of all patients a disparity that cannot be explained by differential longevity alone. ^1, 2^. Despite decades of intensive investigation, the molecular basis for this sex difference remains elusive. Genome-wide association studies have largely identified sex-shared risk loci, and clinical trials have conventionally enrolled pooled cohorts. ^3, 4^. Furthermore, prevailing transcriptomic analyses have typically focused on female-enriched samples or computationally collapsed biological sex into a mere statistical covariate ^5^. Consequently, the current literature is rich in general AD biology but structurally silent on a critical question: whether men and women traverse the same transcriptomic trajectory toward dementia, or if they execute fundamentally divergent molecular programs.

The neurodegenerative transition from normal cognition (NCI) through mild cognitive impairment (MCI) to terminal AD is not a binary event, but a continuous pathophysiological spectrum ^2, 6^. In population-based cohorts such as the Religious Orders Study and Memory and Aging Project (ROSMAP), many specimens occupy intermediate temporal positions in the disease molecular space ^7, 8^. However, the early transition interval particularly from NCI to MCI is notoriously poorly captured by conventional differential expression analysis ^8^. At this pre-clinical stage, individual gene mean shifts are subtle and rarely survive false discovery rate correction. Crucially, this statistical insensitivity does not imply biological latency; rather, it occurs because mean expression tests are mathematically blind to coordinated variance changes that frequently precede mean-level reprogramming ^9, 10^. A transition-aware computational framework capable of tracking variance instability along the NCI-to-AD axis, strictly independent of mean expression, is requisite to unmask early molecular dynamics that standard analyses miss by design.

Here, building upon our recently established geometric framework that models transcriptomic transitions through variance topology, we formally test the hypothesis that the transition to AD is not a unitary molecular cascade quantitatively modified by sex, but rather encompasses two structurally and biologically distinct dynamical architectures. To rigorously evaluate this, we applied our transition-aware framework to bulk RNA-sequencing data from the ROSMAP cohort, strictly stratified by biological sex (n = 401 females, 223 males)

Within this framework, transcripts are not evaluated merely by mean abundance, but are classified into distinct dynamical tiers based on the interval structure of their variance changes ^11^. Specifically, we define a ‘priming’ tier as early, coordinated variance instability emerging at the NCI-to-MCI interval that mathematically precedes measurable mean-level shifts, and a ‘bifurcation’ tier as late-onset variance divergence at the MCI-to-AD threshold, denoting irreversible macroscopic system failure ^11^. For each sex, we classified transcripts into these structural tiers, computed conventional differential expression and pathway enrichment, and mapped both dimensions onto a curated biological process module landscape (BioGraphNode).

By systematically isolating variance instability from mean-level abundance across these stages, our central objective is not merely to catalog which genes change in AD. Rather, we aim to mathematically and biologically resolve whether the temporal structure and underlying module identity of the neurodegenerative trajectory fundamentally diverge between women and men, thereby providing a foundational geometric rationale for sex-stratified mechanistic models and biomarker development.

## Results

### Structurally distinct transcriptomic transition architectures

To determine whether the neurodegenerative trajectory from NCI through MCI to terminal AD follows a shared molecular program across sexes, we initially interrogated the ROSMAP cohort using standard mean-level differential expression gene (DEG) analysis. Although the female cohort is numerically larger across all three diagnostic categories (female *n* = 401, male *n* = 223; **Fig. 1a**), this demographic structure alone cannot account for the profoundly asymmetric transcriptomic response observed between the sexes. As the disease progressed, female AD exhibited a substantially higher DEG burden, particularly culminating in a collapse of 8,935 DEGs during the NCI-to-AD transition, indicating a distinctly mean-shift dominant transcriptomic response. In stark contrast, male AD showed minimal DEG output across all transition pairs (NCI vs. MCI, NCI vs. AD, and MCI vs. AD), yielding a near-total absence of statistically significant mean-level changes (false discovery rate [FDR] < 0.05) (**Fig. 1b**).

**Figure 1.**
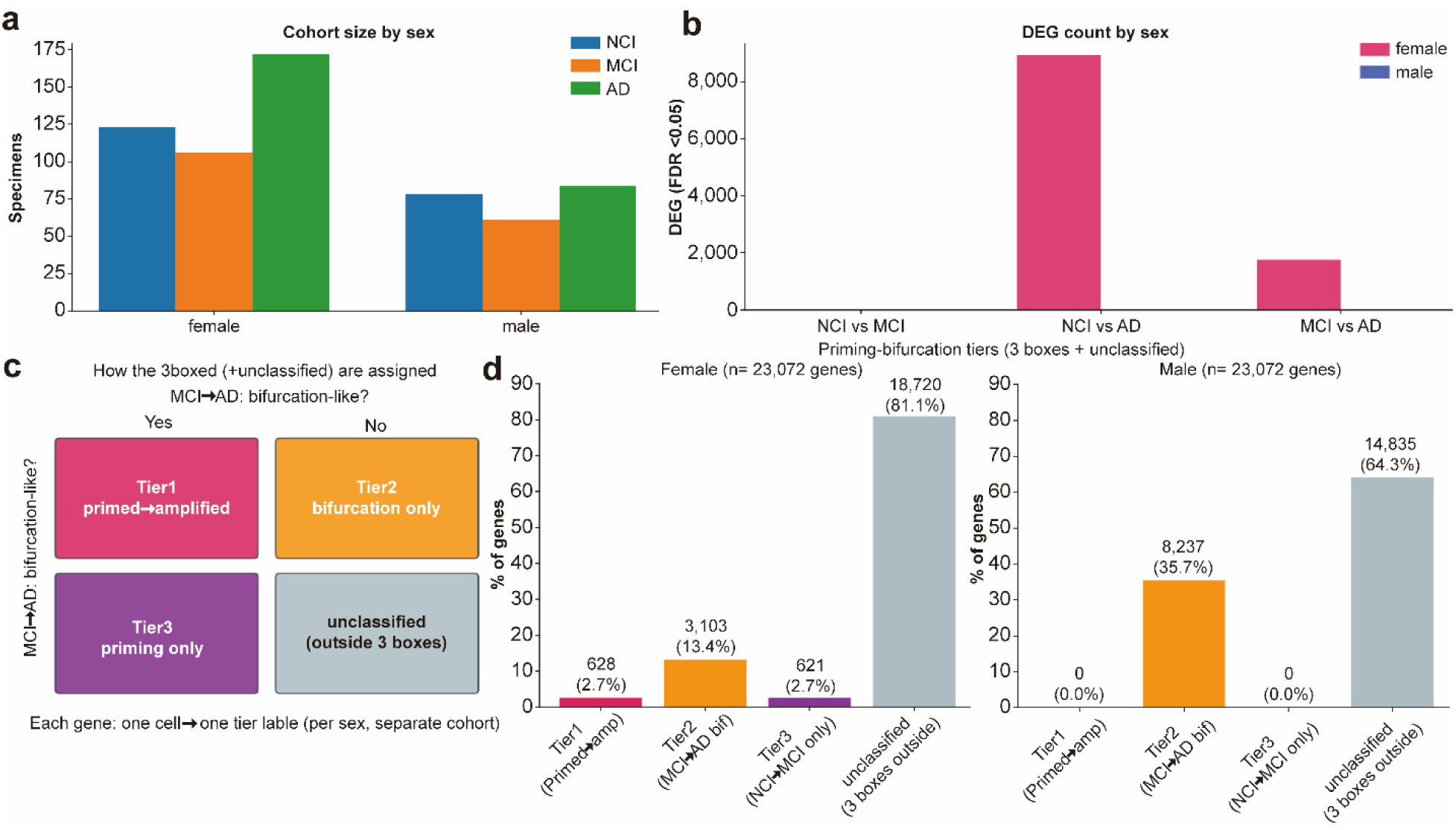
Sex-stratified gene expression architecture reveals fundamentally distinct transcriptomic transition modes in Alzheimer’s disease. (a) ROSMAP cohort composition by sex and cognitive diagnosis. Bar graph showing specimen counts across NCI (cognitively normal), MCI (mild cognitive impairment), and AD diagnosis groups, stratified by female and male cohorts. (b) Differential expression gene (DEG) counts by sex across diagnostic transition pairs (NCI vs. MCI, NCI vs. AD, MCI vs. AD) at FDR < 0.05. (c) Schematic of the three-tier priming-bifurcation classification framework. Each gene receives one tier label per sex, computed in sex-separate cohorts. (d) Bar graph showing the percentage of genes assigned to each priming-bifurcation tier.

To decode these latent transition dynamics, we implemented a ‘priming-bifurcation’ classification framework based on variance instability, which operates strictly independently of mean expression changes. Within this framework, each gene receives one tier label per sex, computed in sex-separate cohorts, and is assigned to one of four categories based on the presence or absence of a priming signal (NCI-to-MCI variance change) and a bifurcation signal (MCI-to-AD variance change): Tier 1 (primed and amplified; both signals), Tier 2 (bifurcation-only; MCI-to-AD variance only), Tier 3 (priming-only; NCI-to-MCI variance only), or unclassified (outside all three distinct boxes) (**Fig. 1c**).

Applying this topological classification to each sex revealed a stark divergence in genome-wide tier distribution (*n* = 23,072 genes). As illustrated by the bar graph showing priming-bifurcation tiers gene percentage, the female cohort showed a predominance of Tier 1 (628 genes, 2.7%) and Tier 2 (3,103 genes, 13.4%), with the majority remaining unclassified (81.1%). Conversely, the male cohort was dominated by Tier 2 bifurcation-only genes (8,237 genes, 35.7%), with absolutely no Tier 1 or Tier 3 priming signals priming signal detected (**Fig. 1d**). This stark male Tier 2 predominance (0 Tier 1) mathematically establishes that male AD progression is governed by a variance-dominant, DEG-silent disease architecture. Importantly, the structural asymmetry of these transition architectures is highly robust. Even under relaxed variance bifurcation thresholds (BIF-relaxed), the stark contrast between the female multi-tier model and the male Tier 2-exclusive collapse is strictly preserved (**Supp. Fig. 1a, b**).

Collectively, these results indicate that the transcriptomic transition in AD is not a unitary molecular cascade quantitatively modified by sex, but rather comprises two structurally independent biological trajectories governed by fundamentally decoupled temporal onsets and dynamical axes.

### Sex-divergent variance acceleration and topological saddle-point localization

To determine the precise physical localization of these distinct transition architectures along the disease continuum, we mapped the transcriptomic variance onto the neuropathological Braak continuum. Mixture modeling and differential covariance (ΔC) metrics consistently identified the MCI stage as a mathematical saddle point within the high-dimensional transcriptomic state space, physically occupying the Braak ∼3.5–4.5 interval. Because these saddle metrics were robustly elevated across both pooled and sex-stratified analyses, they confirm that the MCI stage operates as a universal structural inflection point irrespective of biological sex, even though the peak variance Braak stage diverges between males and females (Fig. 2a).

**Figure 2.**
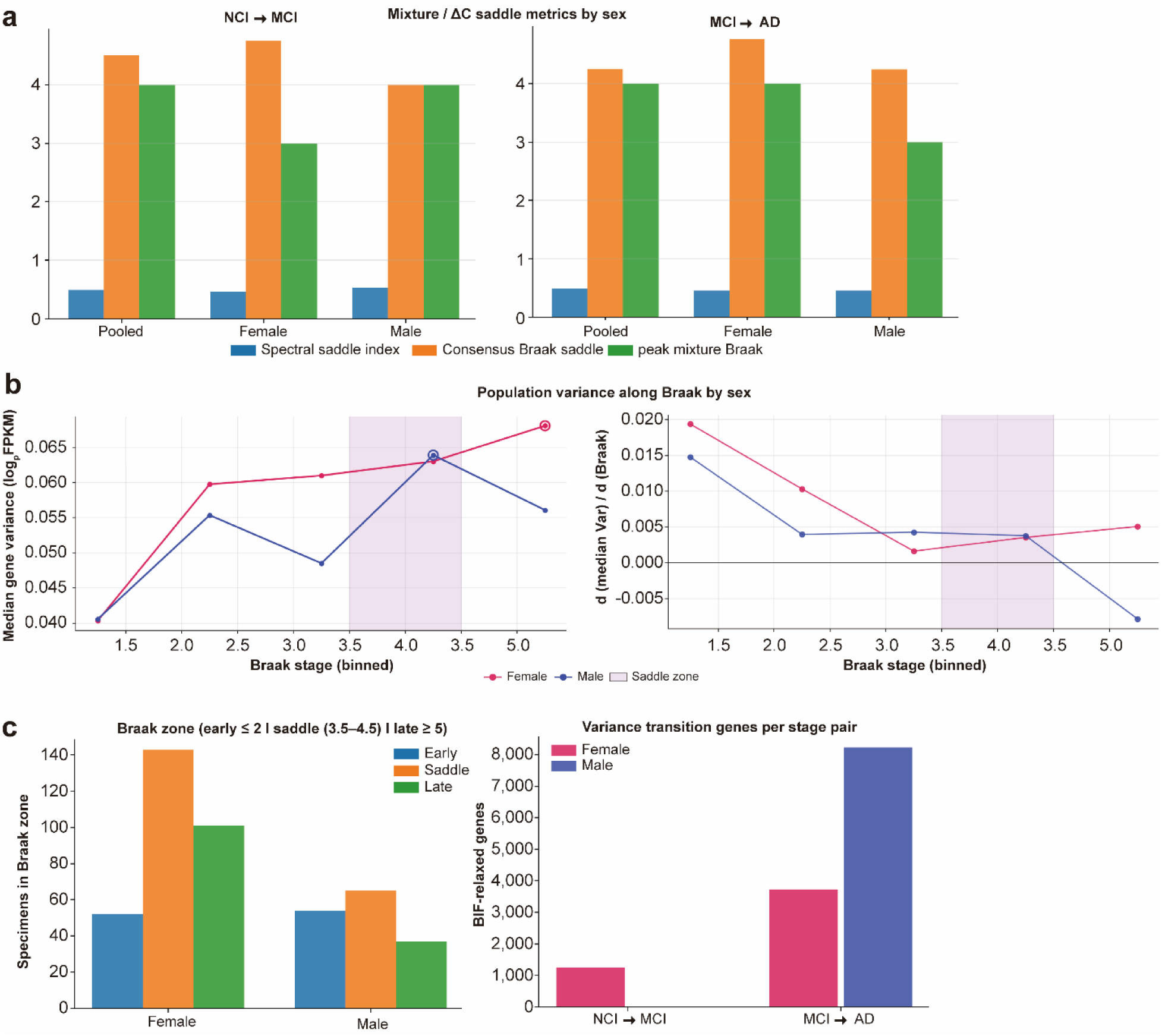
Sex-specific saddle-point localization and transition dynamics along Braak staging. (a) Mixture model ΔC saddle metrics by sex for NCI→MCI (left) and MCI→AD (right) transitions. Three metrics are shown: spectral saddle index (blue), consensus Braak saddle (orange), and peak mixture Braak (green), computed in pooled, female, and male cohorts separately. (b) Median gene expression variance (log10 FPKM) trajectories across Braak stages, stratified by sex (left panel), and their first derivatives (dMedianVar/dBraak; right panel). (c) Sex-stratified transition-point comparison across Braak zones. Left: specimen distribution across early, saddle, and late Braak zones by sex. Center: variance transition gene counts per stage-pair (NCI→MCI, MCI→AD) by sex.

However, the dynamics of variance acceleration through this identified saddle zone (Braak ∼3.5–4.5) fundamentally diverged between females and males. Tracking the median gene expression variance trajectories (log10 FPKM) and their first derivatives (dMedianVar/dBraak) across Braak stages revealed starkly asymmetric geometric profiles. Female transcriptomic variance increases monotonically and peaks in late-stage neuropathology (Braak ≥ 5), with acceleration extending into these late disease stages. Conversely, the male variance trajectory exhibited an earlier, intensely concentrated acceleration peak strictly at the MCI-equivalent Braak window, followed by a subsequent system-wide decline (Fig. 2b).

This temporal dissociation was further corroborated by quantifying specimen distributions across early (≤ 2), saddle (3.5–4.5), and late (≥ 5) Braak zones, alongside gene transition distributions across clinical stages. Evaluating variance-shifting genes per stage-pair clearly contrasted the early NCI-to-MCI priming phenomenon in females with the massive concentration of transition events (6,000–8,000 genes) specifically at the MCI-to-AD interval in males. Furthermore, priming-bifurcation tier counts confirmed that female priming (Tier 1) is prominently concentrated at the NCI-to-MCI interval, whereas the male bifurcation (Tier 2) completely dominates the MCI-to-AD transition (Fig. 2c).

These results indicate that although the MCI stage operates as a universal topological saddle point for both sexes, the temporal dynamics of traversing this critical threshold are strictly sex-divergent: females initiate continuous transcriptomic destabilization well before this clinical boundary, whereas males undergo a catastrophic, system-wide variance collapse exclusively upon crossing it.

### Orthogonal mapping of immune variance priming and synaptic mean-level collapse in females

To determine whether the early pre-clinical variance instability and the late-stage mean-level collapse observed in the female cohort represent sequential steps of a single pathological cascade or operate as independent biological programs, we mapped the functional ontology of both the mean-shifted and variance-destabilized gene pools. Conventional pathway enrichment of the massive pool of DEGs downregulated across the longitudinal NCI-to-AD transition revealed a dominant concentration in neuroactive ligand-receptor interactions, G protein-coupled receptor (GPCR) signaling, and neuropeptide pathways (**Fig. 3a**). Furthermore, examining the NCI-to-AD top log2-fold change gene set (n ≤ 400) explicitly mirrored this GPCR and olfactory receptor axis, confirming profound chemosensory and neuromodulatory gene suppression. Conversely, upregulated DEGs in the NCI-to-AD transition were primarily enriched for extracellular membrane and cell surface components (GO:CC), and the specific MCI-to-AD transition pattern further highlighted a late-stage synaptic loss signature. This network profile constitutes a coherent synaptic and neuromodulatory failure signature that classically drives the cognitive collapse in terminal AD.

**Figure 3.**
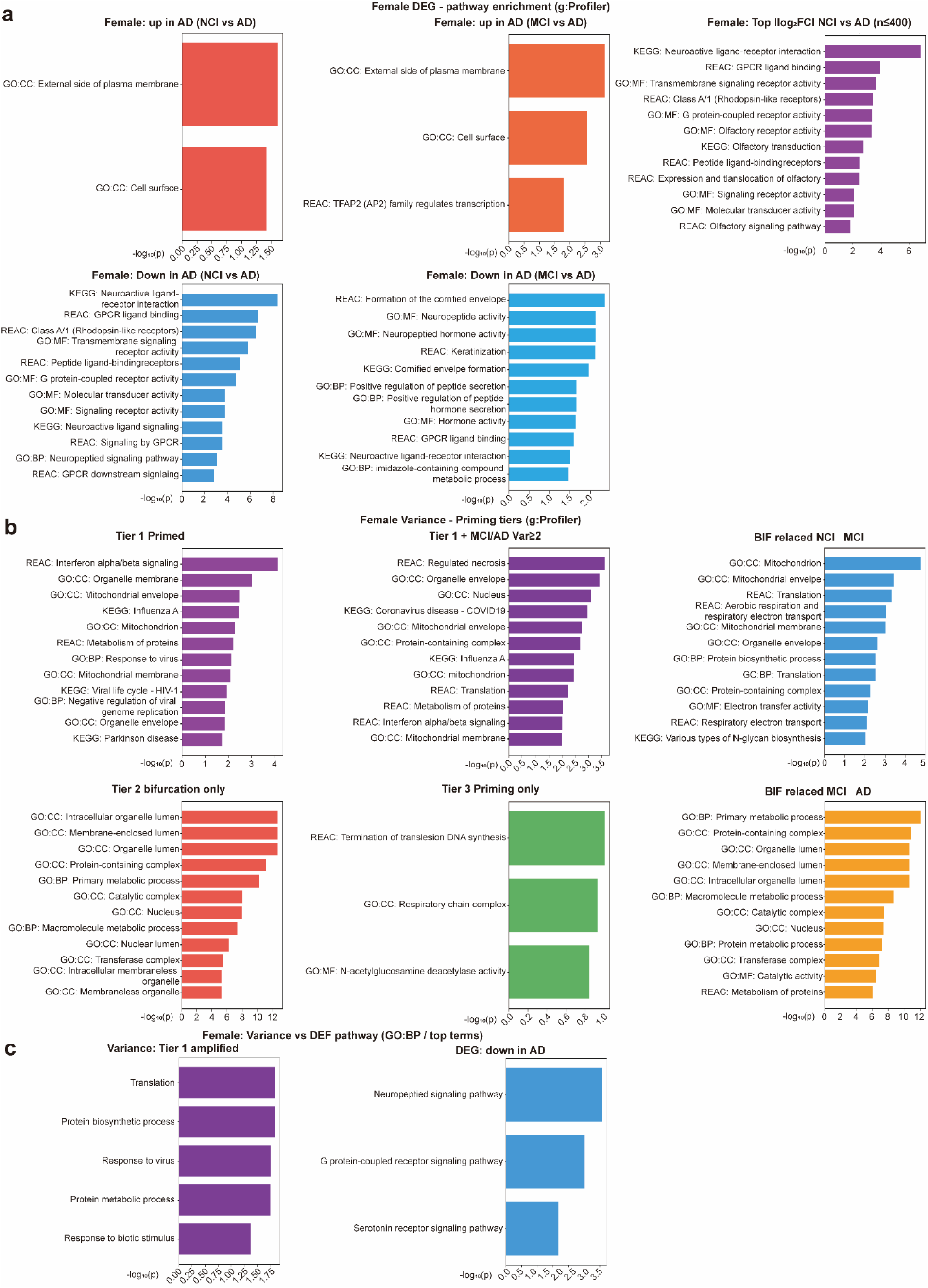
Female AD transcriptomics: pathway enrichment of DEG and variance tiers reveals dual-axis disease mechanism. (a) g:Profiler pathway enrichment analysis of female-specific DEGs across diagnostic comparisons (NCI vs. AD, MCI vs. AD). The analysis includes upregulated and downregulated gene sets, as well as the NCI vs. AD top log2FC gene set (n ≤ 400), mapped across GO:CC, KEGG, and Reactome (REAC) databases. (b) g:Profiler pathway enrichment analysis of female variance tiers (Tier 1, Tier 2, and Tier 3) using REAC and GO databases. The analysis includes additional conditions evaluating Tier 1 combined with MCI/AD variance-squared metrics and BIF-relaxed transition sets (NCI→MCI and MCI→AD). (c) Comparative pathway mapping between female Tier 1 (variance-amplified) and DEG-downregulated gene sets, specifically evaluated at the Gene Ontology Biological Process (GO:BP) level.

However, the early molecular drivers of the female transition operate on an entirely different analytical axis. The 628 Tier 1 genes, which exhibit premature variance instability specifically at the pre-clinical NCI-to-MCI interval, are functionally distinct from the terminal synaptic DEG pool. Instead, this early variance-amplified network is highly enriched for innate immune activation, particularly interferon (IFN) alpha/beta signaling and viral response pathways, alongside organelle membrane and mitochondrial processes (REAC/GO) (**Fig. 3b**). Expanding this analysis, combining Tier 1 with MCI/AD variance-squared metrics further amplified these immune and mitochondrial terms while adding translation and metabolism signatures. Other variance classifications also showed distinct localizations: Tier 2 (bifurcation-only) enriched for intracellular organelle and lumen terms, while Tier 3 (priming-only) captured respiratory chain complex enrichment. Importantly, BIF-relaxed sets (NCI-to-MCI and MCI-to-AD) confirmed that organelle membrane and mitochondria act as sustained variance amplification hubs across the transition.

The strict orthogonality of these two functional axes is directly visualized through comparative pathway mapping at the GO:BP (Biological Process) level. This analysis explicitly demonstrated that Tier 1 amplified genes are highly enriched for translation, protein biosynthesis, response to virus, and protein metabolic processes, whereas DEG-downregulated genes are restricted to neuropeptide signaling, GPCR signaling, and serotonin receptor pathways (**Fig. 3c**). This topological comparison confirmed that the early immune variance network and the subsequent synaptic mean-collapse network do not substantially overlap.

Collectively, these results indicate that innate immune priming and terminal synaptic failure in females are not parallel reflections of a single disease cascade observed through different statistical lenses; rather, they represent two structurally distinct biological processes—specifically, early translational and proteostatic stress operating entirely independently from downstream synaptic and neuromodulatory loss—that drive the female AD trajectory through fundamentally decoupled analytical spaces and temporal windows.

### Direct visualization of mean–variance axis dissociation

To definitively validate whether the orthogonal biological architectures identified through pathway mapping manifest as fundamentally distinct transcriptional behaviors at the individual patient level, we juxtaposed single-specimen expression heatmaps of DEG-selected versus variance-tier-selected gene sets (top-50 ranked genes per sex). The female top DEG panel demonstrated a coherent, graded, and monotonic mean-level transition across NCI→MCI→AD, consistent with a linear priming-and-amplification model (**Fig. 4a, left**). Conversely, the male top DEG panel showed a more heterogeneous expression pattern with a markedly lower overall mean-shift magnitude, consistent with the severely reduced DEG count in males (**Fig. 4a, right**) and indicative of an MCI-silent, AD-onset mean-shift model even within the DEG subset (**Supp. Fig. 3a**).

**Figure 4.**
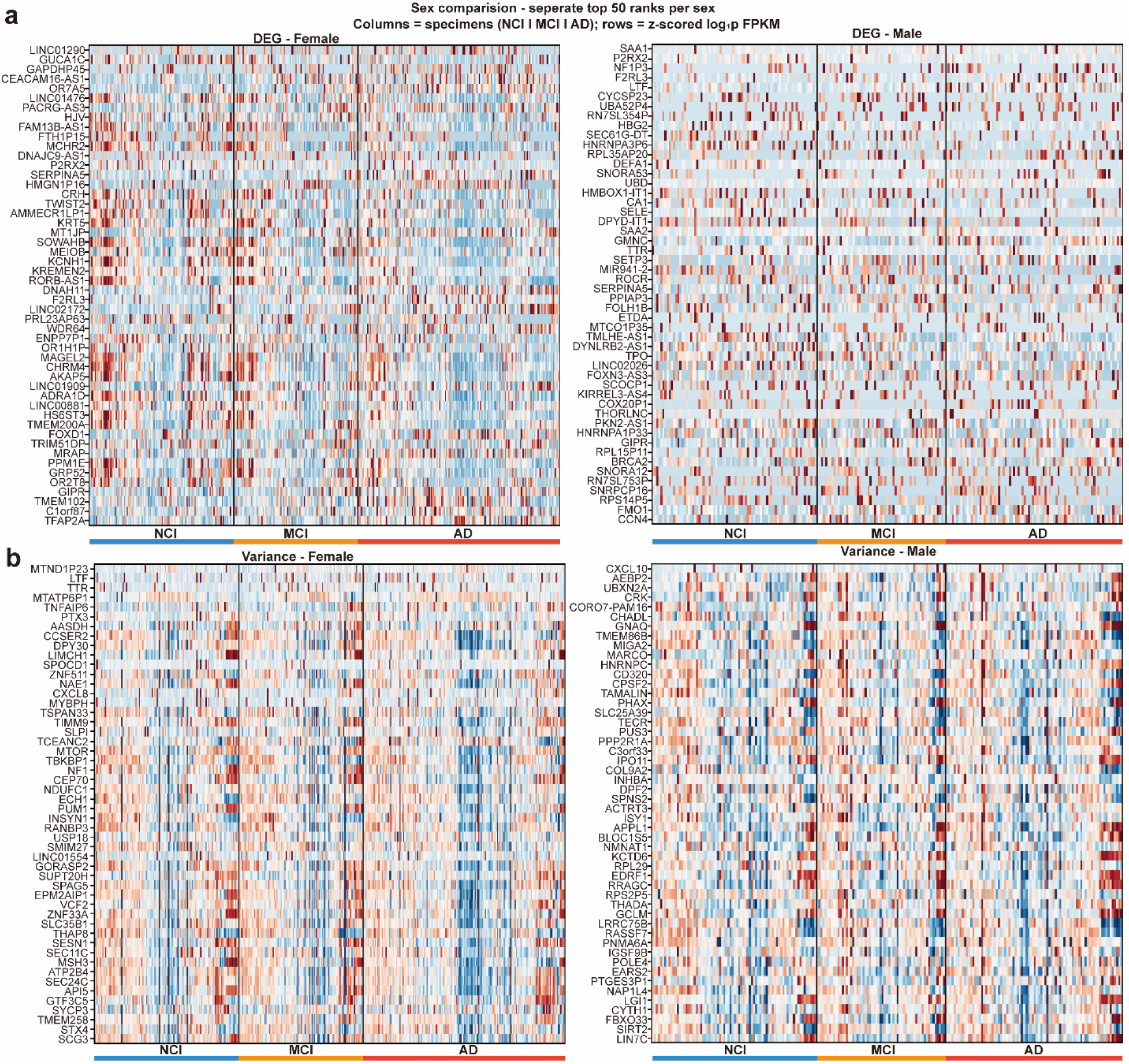
Sex-specific gene expression heatmaps reveal distinct signal architectures for DEG and variance top-ranked genes. (a,b) Heatmaps showing z-scored log1p FPKM expression across individual specimens (columns: NCI | MCI | AD, indicated by the bottom color bars) for the top-50 ranked genes per sex. Each gene set is ranked independently within sex. (a) Heatmaps of the top-50 DEG-ranked genes for the female cohort (left) and the male cohort (right). (b) Heatmaps of the top-50 variance/BIF-ranked genes for the female cohort (left) and the male cohort (right).

In sharp contrast, the variance-ranked panels revealed distinct signal architectures invisible to mean-based analysis. The female variance (Tier 1) panels exhibited extreme population heterogeneity and mixed-direction expression patterns without monotonic progression, visually confirming that early immune priming operates exclusively as a variance phenomenon rather than a structured mean-level shift (**Fig. 4b, left**). Similarly, the male variance (Tier 2) panels displayed a massive bifurcation-like pattern without any consistent monotonic mean separation across the diagnostic groups; specifically, subsets of specimens within the same diagnostic stage showed highly divergent expression values (dark red vs. dark blue) (**Fig. 4b, right; Supp. Fig. 3a, b**). This bimodal distribution within the MCI and AD columns directly reflects the population-splitting signature predicted by the kappa-manifold bifurcation model.

Further aggregation into group-mean heatmaps firmly corroborated these individual-level dynamics. Male variance top genes—including key immune-regulatory loci (PTX3, CXCL10) and membrane trafficking genes (BNIP3, VAMP family)—exhibited near-zero mean shifts at the NCI-to-MCI interval, followed by striking AD-stage bifurcation (**Supp. Fig. 3b**). This profound lack of group-mean divergence provides compelling visual evidence that male AD biology cannot be reliably captured through classical mean-expression analyses, further validating MCI as a definitive mathematical saddle point.

Collectively, these visual representations indicate that the neurodegenerative transition is not governed solely by uniform, mean-level functional decline; rather, both the early innate immune priming in females and the massive late-stage disruption in males operate through profound, population-splitting variance bifurcations that constitute the true dynamical forces of disease progression.

### System-wide organelle membrane variance coupling drives the male transition

Having established through visual expression mapping that the male AD trajectory is exclusively driven by a massive, mean-silent variance bifurcation, we sought to decode the precise biological identity of this 8,237-gene Tier 2 pool by projecting it onto a three-dimensional BioGraphNode topological landscape. This funnel architecture enables topology-aware pathway analysis independent of flat enrichment cutoffs by assigning genes from the full transcriptome through GOA annotation layers to specific functional modules. Unlike the multi-tiered female trajectory, the male transition program collapses entirely into this single block that bifurcates strictly at the late MCI-to-AD interval. The topological projection revealed that the male Tier 2 architecture manifests as a highly active, broad “hot cloud” spanning membrane, trafficking, and post-translational modification (PTM) spaces. This starkly contrasts with the comparatively narrow footprint of the female Tier 2 program (**Fig. 5a**).

**Figure 5.**
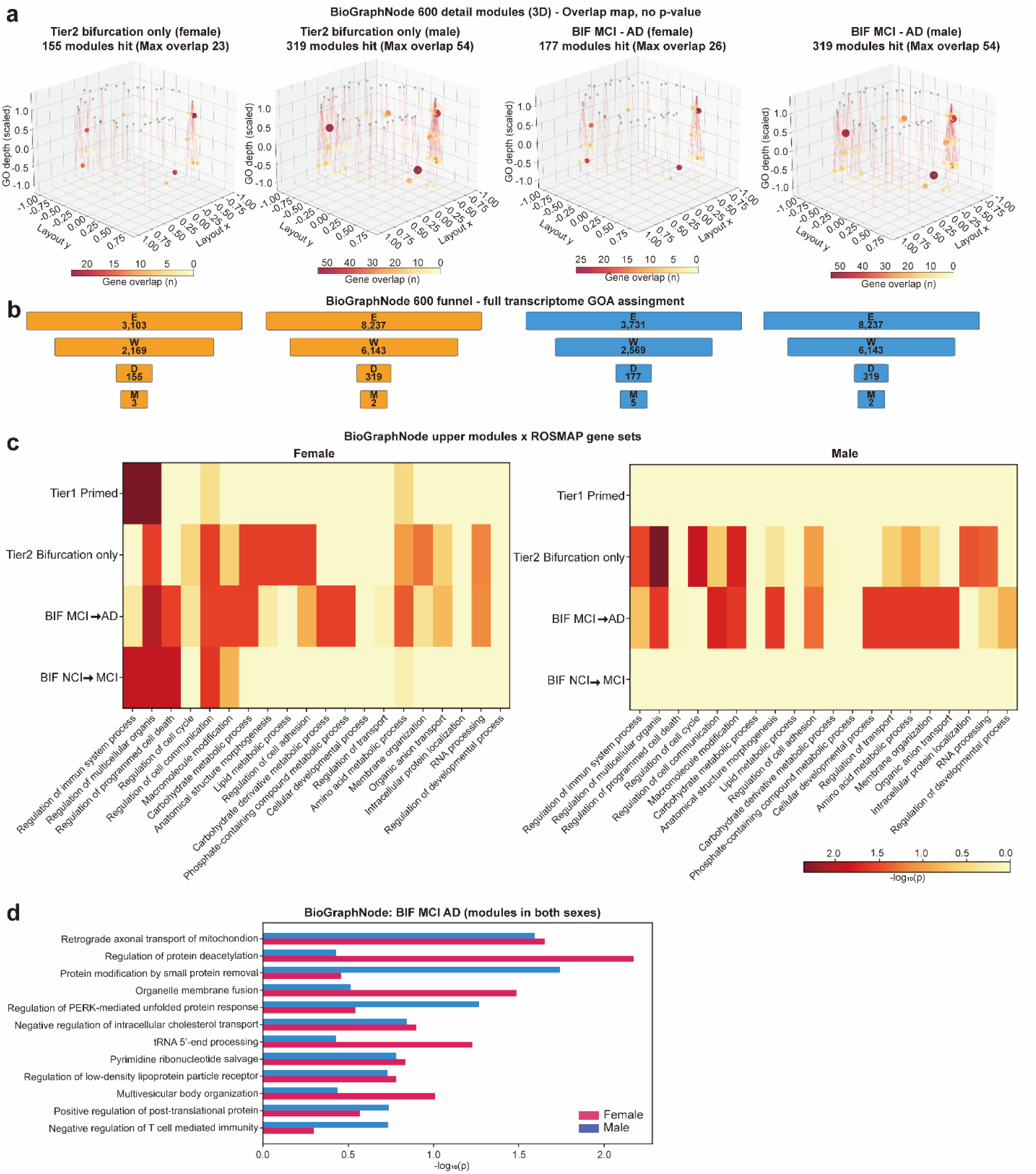
BioGraphNode network topology maps sex-divergent AD transition modules onto biological process space. (a) Three-dimensional BioGraphNode funnel scatter plots for female Tier 2 (NCI→MCI), female BIF MCI→AD, male Tier 2 (NCI→MCI), and male BIF MCI→AD (left to right). Each bubble represents a GOA (Gene Ontology Annotation) module; bubble size reflects the number of overlapping genes; color encodes enrichment magnitude. (b) BioGraphNode gene set funnel statistics. For each sex-transition combination, cascading boxes show: total Ensembl gene count (E), GOA-annotated subset (W), directionally-assigned module genes (D), and multi-assigned genes (M). (c) Cross-sex BioGraphNode module enrichment heatmaps. Rows represent variance/priming tier categories (Tier1_primed, Tier2_bifurcation_only, BIF_MCI_AD, BIF_NCI_MCI); columns represent GOA biological process modules. Color encodes -log_10_(p). (d) Divergent bar chart displaying shared BIF_MCI_AD modules enriched in both cohorts (female: pink; male: blue).

Funnel statistics successfully mapped this extensive gene pool (total Ensembl gene count, E = 8,237) to 319 highly specific functional directionally-assigned modules (D = 319) from a GOA-annotated subset (W = 6,143). In contrast, female Tier 2 and BIF-MCI→AD subsets retained proportionally lower module assignment rates, consistent with their smaller gene set sizes (**Fig. 5b and Supp. Fig. 4a**). Cross-sex module enrichment heatmaps clarified that while females show prominent enrichment in immune, cell cycle, and signal transduction modules, classical neuroinflammation is not the primary driver in males; rather, the male signal is heavily concentrated within the organelle membrane and vesicular trafficking axes, alongside carbohydrate/lipid metabolism and RNA-processing modules (**Fig. 5c and Table 1**).

**Table 1.** Top BioGraphNode overlap modules, male Tier2/BIF (n = 8,237 genes; 6,143 with GOA annotation).

| GO:BP module | n_overlap | Mechanistic context |
| --- | --- | --- |
| Organelle membrane fusion | 54 | SNARE complex dynamics; vesicle docking |
| Protein modification by small protein removal | 50 | Ubiquitin/SUMO deconjugation |
| Post-translational protein modification (broad) | 45 | Proteostasis maintenance landscape |
| Gliogenesis | 30 | Glial biology; potentially reactive gliosis |

This structural mapping is fully corroborated by standard pathway analyses, which similarly highlight extensive reorganizations in membrane trafficking, mitochondrial, and endosomal systems (**Supp. Fig. 2b, c**). Nevertheless, analysis of the bifurcation modules shared across both sexes identified a specific set of convergent pathways. These included retrograde axonal transport of mitochondria, organelle membrane fusion, regulation of low-density lipoprotein receptor, negative regulation of intracellular cholesterol transport, protein modification by small protein removal, and most notably, the PERK-mediated unfolded protein response (UPR) and multivesicular body (MVB) organization pathways. This critical intersection suggests that the male-specific membrane trafficking bifurcation ultimately converges with a global proteostatic failure axis observed in both sexes (**Fig. 5d**).

Collectively, these results indicate that male AD progression represents a fundamental, variance-dominant disruption of intracellular membrane dynamics, which, despite operating through an entirely divergent biological axis from the female program, ultimately converges into a universal, terminal collapse of global proteostasis.

## Discussion

The central finding of this study establishes that the transcriptomic transition to AD does not follow a unified molecular program that is merely quantitatively modified by biological sex. Instead, female and male brains within the ROSMAP cohort exhibit qualitatively distinct transition architectures that fundamentally differ in their temporal onset, the biological networks engaged, and the analytical axis (mean versus variance) on which the dominant signals reside.

In females, the transition is structured and sequential: an early IFN and immune variance priming phase is detectable at the NCI-to-MCI interval, bifurcation expands toward MCI-to-AD, and which is subsequently followed by a large-scale, mean-level synaptic collapse during terminal AD progression. These events are strictly temporally ordered and occupy completely distinct biological spaces. Conversely, the male transition is concentrated and delayed: a singular, massive Tier 2 variance block bifurcates at the MCI-to-AD threshold across organelle membrane biology, exhibiting an absolute absence of early immune priming or detectable mean-level expression changes. These two programs are not merely mirror images; they engage entirely different molecular machineries at distinct disease stages through divergent expression dynamics.

This structural distinction carries direct implications for interpreting the sex disparity in AD incidence ^3^. The female excess in AD prevalence is not simply a secondary consequence of differential longevity. The female transition program initiates significantly earlier at the pre-clinical NCI-to-MCI phase with an immune priming wave and generates a massive synaptic collapse that is entirely absent in the male cohort. Importantly, male biology is not silent; rather, it operates on an orthogonal variance-dominant channel that conventional DEG analyses are mathematically ill-equipped to resolve.

The female Tier 1 IFN and immune priming signature (628 genes), heavily enriched for IFNα/β signaling and viral response, represents the earliest sex-specific molecular event in this dataset. This priming is detectable precisely at the NCI-to-MCI boundary, preceding any measurable mean-level shifts. The subsequent AD mean-level functional decline characterized by the loss of synaptic, GPCR, and neuropeptide pathways (8,935 FDR-significant DEGs) is strictly temporally downstream and biologically orthogonal, sharing no significant gene identity with the early Tier 1 priming pool. This two-wave architecture suggests that innate immune and interferon activation operating entirely at the variance level to drive population-wide transcriptional destabilization may act as an upstream instigator that precipitates the downstream synaptic vulnerability culminating in terminal mean-level collapse.

While this hypothesis aligns with emerging literature implicating type I interferon signaling in early neuroinflammatory priming, prior investigations have consistently failed to resolve the temporal relationship between immune variance activation and synaptic mean-level failure within the same analytical scale. However, we explicitly acknowledge that the cross-sectional design of the ROSMAP cohort precludes direct causal inference. The sequential interpretation of Tier 1 priming preceding Tier 2 bifurcation and downstream DEG loss remains a population-level structural inference requiring rigorous validation in longitudinal or pseudo-time frameworks.

The male transcriptomic transition is dominated by a massive Tier 2 bifurcation block comprising 8,237 genes at the MCI-to-AD threshold, comprehensively mapped to 319 specific BioGraphNode functional modules ^11, 12^. This architectural collapse is emphatically not a statistical artifact of sample size. A randomized gene set of equivalent magnitude would yield diffuse and incoherent module overlaps; instead, the observed male signal demonstrates profound biological specificity, overwhelmingly concentrated in organelle membrane fusion (54 overlapping genes), SNARE-complex dynamics, ubiquitin/SUMO deconjugation, and proteostasis modules reflects genuine biological specificity.

The mechanistic interpretation of this architecture is that the male AD transition is driven by a system-wide variance coupling across the intracellular membrane trafficking machinery. This explicitly includes the SNARE-complex dynamics governing vesicle docking, the ubiquitin/SUMO pathways regulating organelle membrane identity, and the associated global proteostatic landscape. Crucially, this system-wide destabilization is mechanistically highly coherent with the established roles of endo-lysosomal and membrane trafficking networks in APP processing and tau secretion, physically bridging our variance-based transcriptomic signatures with classical AD proteinopathies ^13, 14^.

Crucially, the absence of statistically powered DEGs in the male cohort must not be misinterpreted as an absence of pathological biology. It strictly represents an absence of detectable mean-level reprogramming at the current sample size (n = 223). The underlying variance signal is massive, coherent, and highly structured. Male AD biology in this cohort is fundamentally variance-dominant. While larger male cohorts may eventually yield mean-level shifts, such future findings will not negate the foundational discovery that the primary, most accessible disease signal in males operates precisely on the variance axis.

The identification of 1,249 variance bifurcation events at the NCI-to-MCI interval exclusively in females redefines MCI not merely as a clinical diagnosis, but as a strictly sex-divergent molecular inflection point. In the female trajectory, clinical MCI is already accompanied by measurable, transcriptome-wide variance divergence; conversely, in the male trajectory, MCI represents a structurally quiet latency period immediately preceding late-stage catastrophic bifurcation. This profound asymmetry carries immediate clinical significance. Biomarkers designed to assess female AD risk at the MCI stage must strategically target variance-level immune priming rather than mean-level synaptic loss, which has not yet manifested at this pre-clinical stage. Conversely, male MCI biomarkers must be variance-based tools explicitly designed to detect the impending destabilization of organelle membrane trafficking machinery. Current clinical paradigms treat MCI staging equivalently across sexes ^1, 6^; our molecular evidence strongly indicates that this clinical equivalence is biologically unjustified.

The findings presented herein argue definitively against pooled transcriptomic analyses in AD transition cohorts. Arithmetically merging these two distinct biological programs fundamentally degrades signal integrity: it systematically suppresses the female variance priming signal (which is too structurally specific to survive cohort averaging), dilutes the male organelle membrane signature (which is concentrated within a massive but mean-flat pool), and ultimately generates an underpowered, hybrid DEG profile that accurately represents the biology of neither sex.

For successful clinical translation, sex-stratified biomarker discovery is no longer a secondary consideration it is an absolute prerequisite for diagnostic accuracy. An IFN and immune variance signature at the NCI-to-MCI transition offers high utility as a female-specific early risk indicator, whereas perturbations in organelle membrane trafficking constitute a highly specific male MCI-to-AD transition marker. Furthermore, these distinct transcriptomic programs nominate partially divergent therapeutic targets. Interventions focused on synaptic maintenance may yield disproportionate efficacy in female populations, whereas therapeutics aimed at stabilizing vesicular trafficking and proteostasis may be critical for halting male disease progression. Clinical trial designs that ignore this fundamental structural distinction risk systematically masking real, yet entirely sex-confined, efficacy signals.

This study acknowledges several methodological constraints that warrant consideration. Firstly, the male cohort size (*n* = 223) renders conventional mean-level DEG analysis statistically underpowered, making male mean-axis interpretations explicitly exploratory, although the variance-axis signal remains mathematically robust. Secondly, the reliance on bulk DLPFC RNA-seq inherently conflates purely cell-autonomous intracellular transcriptional reprogramming with macro-level shifts in cellular composition across the AD brain. Thirdly, the cross-sectional nature of the cohort establishes a population-level variance structure but cannot definitively confirm the longitudinal sequence of events within individual patients, meaning the sequential progression model (priming to bifurcation to mean collapse) remains a structural hypothesis. Furthermore, our analytical focus on the transition corridor (Braak ∼3.5–4.5) implies that earlier, highly localized molecular events at lower Braak stages might remain undetected within the current cohort structure. Finally, regarding BioGraphNode overlap metrics, standard enrichment p-values approach zero at extreme gene set sizes (e.g., 8,237 genes); thus, module overlap counts are strictly utilized as topological landscape descriptors, with biological interpretations grounded entirely in mechanistic coherence rather than statistical significance alone.

To advance these findings, priority future directions include executing sex-stratified single-nucleus RNA-seq in transition-enriched cohorts to resolve variance tier architectures at the definitive cell-type level. Given the terminal nature of human brain tissue sampling, validating the proposed priming-to-bifurcation temporal ordering requires dual strategies: deploying advanced pseudo-time computational frameworks on single-cell datasets to infer temporal dynamics within the brain, and utilizing true longitudinal transcriptomic profiling of accessible peripheral biofluids (e.g., CSF or blood) to track within-individual progression. Mechanistic functional validation should involve targeted follow-ups of the organelle membrane fusion module, specifically SNARE-complex regulators identified in the male Tier 2 network, utilizing sex-stratified cellular and in vivo models. Furthermore, integrating existing ROSMAP amyloid and tau quantification metrics with individual tier memberships will definitively test whether variance bifurcations temporally precede, or merely co-occur with, measurable macroscopic proteinopathies. Finally, performing rigorous power analyses across projected, larger-scale male cohorts is necessary to establish whether male mean-level biology eventually converges with, or remains strictly orthogonal to, the massive variance signals identified in this study.

## Conclusion

This study fundamentally redefines the molecular landscape of AD progression by shifting the analytical paradigm from classical mean-based differential expression to second-order variance and co-variance dynamics. Crucially, we demonstrate that covariance-based transcriptomic analysis enables the precise identification of early-stage network destabilizations significantly before these disruptions can be detected by standard mean-level DEG analysis. Furthermore, our findings establish that AD does not follow a unitary trajectory but progresses through completely divergent, sex-specific biological programs. The female transition is initiated by an early, variance-amplified immune priming phase driven primarily by IFN signaling. In stark contrast, the male trajectory operates on an entirely different channel, characterized by a massive, mean-silent structural collapse across organelle membrane and vesicular trafficking networks. Ultimately, the discovery of these orthogonal disease architectures mandates an immediate paradigm shift in clinical translation. Recognizing that men and women undergo fundamentally different paths of disease development highlights the absolute necessity of adopting sex-specific precision medicine approaches for both early diagnostic biomarker detection and targeted therapeutic intervention in AD.

## Methods

### Data Source and Acquisition

Transcriptomic profiles were obtained from the dorsolateral prefrontal cortex (DLPFC) of participants in ROSMAP. ^15, 16^. To rigorously interrogate sex-specific molecular trajectories, the cohort was strictly partitioned into female (n = 401) and male (n = 223) subgroups prior to any analytical processing. The disease transition corridor was defined using standardized clinical cognitive diagnoses, spanning NCI, MCI, and AD. All subsequent statistical evaluations and functional mappings were executed independently for each sex, ensuring that distinct biological programs were not arithmetically obscured by sample pooling.

### Data Preprocessing and Normalization

To establish a robust quantitative foundation and minimize the influence of technical noise, raw transcriptomic profiles were systematically processed prior to variance and mean-level evaluations ^11^. Expression abundances, initially quantified as Fragments Per Kilobase Million (FPKM), were subjected to a log-transformation specifically, log_2_(FPKM+1) to stabilize systemic variance and strictly mitigate inherent heteroscedasticity across the high-dimensional dataset ^17, 18^. We subsequently implemented a stringent low-expression exclusion filter to isolate biologically active transcripts from stochastic background noise. Transcripts demonstrating zero detectable expression in over 80% of the aggregate cohort were categorically removed from the feature space. This rigorous quality control protocol yielded a high-confidence, transcriptomically active universe comprising precisely 23,072 genes. Crucially, this standardized 23,072-gene matrix served as the identical mathematical background for all downstream sex-stratified tier classifications and pathway enrichments, ensuring that the fundamentally divergent female and male topological architectures discovered in this study are true biological phenomena, rather than artificial byproducts of differential baseline gene exclusion.

### Transition-Aware Tier Classification

Building upon our previously established bifurcation-intermediate variance analytical framework, we implemented a variance-centric classification strategy to capture transcriptomic network instability independently of conventional mean-level expression shifts ^11^. By applying this validated Riemannian and variance-based approach to the sex-stratified cohorts, we aimed to isolate sex-specific topological phase transitions that remain inherently invisible to classical mean-based models.

Genes were evaluated across the sequential NCI→MCI→AD clinical continuum and categorized into four structural tiers based on their geometric variance trajectories:

- **Tier 1 (Primed-Amplified):** Encapsulates genes exhibiting primed variance at the initial NCI→MCI interval followed by amplified variance during the MCI→AD progression. Biologically, this tier represents an early, coordinated network instability that systematically escalates acting as a mechanistic ‘priming’ foundation for subsequent disease progression.
- **Tier 2 (Bifurcation-Only):** Comprises genes displaying isolated variance bifurcation exclusively at the late MCI→AD transition, notably without any prior Tier 1 priming. This tier reflects a late-onset variance divergence and mathematically constitutes the overwhelmingly dominant transcriptomic transition signal within the male trajectory.
- **Tier 3 (Priming-Only):** Represents transient variance inflation strictly localized to the initial NCI→MCI boundary, completely lacking subsequent MCI→AD amplification. This delineates an early molecular instability that does not escalate further into terminal disease stages.
- **Unclassified:** Denotes transcripts that do not satisfy these strict variance threshold rules. Importantly, these genes are not transcriptomically inert; rather, they map to the broader functional module background, serving as the foundational landscape that supports the global topological manifold architecture.

This established tier classification enabled the direct geometric decoupling of early-stage immune priming from late-stage structural collapse within each biological sex.

### Differential Expression and Pathway Enrichment

To contextualize our variance-centric findings against classical paradigms, standard mean-level DEG evaluations were performed across all pairwise diagnostic intervals (NCI vs. MCI, NCI vs. AD, and MCI vs. AD). These analyses were executed independently per sex utilizing a standardized limma/DESeq2-based computational pipeline, establishing statistical significance at a strict False Discovery Rate (FDR) of < 0.05 ^17, 19^. Functional mapping of these mean-shifted signatures was conducted using the g:Profiler suite ^20^, querying established functional ontologies (e.g., GO:BP, KEGG, and Reactome) ^12, 21, 22^. To guarantee stringent statistical specificity, all enrichment analyses explicitly utilized the defined 23,072-gene universe as the computational background, applying an enrichment threshold of FDR < 0.05. Crucially, we acknowledge a fundamental statistical constraint regarding the male cohort (n = 223). At this specific sample size, bulk transcriptomic DEG analysis across heterogeneous cortical tissue is systematically underpowered to detect subtle mean-level pathological shifts. Consequently, all male mean-level pathway enrichments are explicitly designated as exploratory within this study. This formal designation mathematically prevents the critical misinterpretation of an absent scalar signal as a true absence of underlying biological pathology—a foundational limitation that our variance-based topological framework specifically overcomes.

### BioGraphNode Module Overlap

To resolve the functional architecture of the sex-specific transition programs, we developed and deployed a customized BioGraphNode compendium. This hierarchical framework integrates approximately 600 highly resolved Gene Ontology Biological Process (GO:BP) detail modules and 48 upper-level rollup categories ^12^, comprehensively mapped against the established whole-transcriptome background of roughly 23,000 genes. Each sex- and tier-stratified gene set was systematically projected onto this compendium to compute absolute module overlap counts (n_overlap_).

We explicitly acknowledge a fundamental statistical constraint regarding hyper-geometric enrichment evaluations: for extraordinarily large gene pools—most notably the male Tier 2 network encompassing n = 8,237 transcripts—standard enrichment *p*-values arithmetically converge to zero for virtually any broad biological module ^23^.

Consequently, rather than relying on mathematically inflated probability metrics, absolute module overlap counts were rigorously utilized as network landscape descriptors to ascertain functional concentration and mechanistic coherence. To fully capture the systemic structural divergence between the sexes, these architectural distributions were embedded and visualized within a three-dimensional force-directed graph environment ^24^, where node parameters encode n_overlap_, thereby translating discrete gene networks into a measurable functional landscape.

## Data availability

The primary human transcriptomic and clinical-pathological datasets analyzed in this study were obtained from the Religious Orders Study and Rush Memory and Aging Project (ROSMAP), managed by the Rush Alzheimer’s Disease Center, Chicago. These data are publicly accessible via the AD Knowledge Portal (https://adknowledgeportal.org) under Synapse ID: syn3219045. All intermediate processed differential covariance matrices and generated secondary statistical data supporting the findings of this study are available within the article, or from the corresponding author (Do-Geun Kim) upon reasonable request.

## Code availability

The mathematical pipelines and custom algorithms developed for the geometric trajectory analysis, differential covariance (ΔC) spectral decomposition, and Riemannian manifold mapping are mathematically detailed in the Methods. The core computation—encompassing stage-wise sample covariance matrix formulation, shrinkage estimation, and Log-Euclidean metric projection—was executed via an in-house analytical platform developed by Do-Geun Kim’s laboratory. Due to ongoing intellectual property protections regarding the core tensor scaling algorithms, the custom software framework and associated scripts are not publicly deposited but are available from the corresponding author upon reasonable request. Key standard software and packages utilized include R (v4.2.1) and Python (v3.9) for core data processing, limma and DESeq2 for mean-level differential expression analysis, mclust (v5.4.1) for mixture modeling, SciPy (v1.10.0) for von Neumann entropy computation, g:Profiler (gprofiler2 v0.2.2) for functional enrichment analysis, and for three-dimensional force-directed topological manifold visualization.

## Acknowledgements

We express our deepest gratitude to the participants of the Religious Orders Study and the Rush Memory and Aging Project for their invaluable contributions of data and post-mortem tissue, which made this dynamic network modeling possible.

## Contributions

S.B., and D-G.K. conceived and designed the study, developed the methodology and analysis pipeline, and carried out the formal data analysis, encompassing bioinformatics, statistical evaluation, and modeling. S.B., and D-G.K. provided resources, secured funding, supervised the project, and managed project administration. M.C. and S.B., and D-G.K. wrote the original draft of the manuscript, performed the visualization of figures and data plots, and contributed to reviewing and editing the manuscript.

## Ethics declarations / Competing interests

The authors declare no competing interests.

